# Evidence-based calibration of computational tools for missense variant pathogenicity classification and ClinGen recommendations for clinical use of PP3/BP4 criteria

**DOI:** 10.1101/2022.03.17.484479

**Authors:** Vikas Pejaver, Alicia B. Byrne, Bing-Jian Feng, Kymberleigh A. Pagel, Sean D. Mooney, Rachel Karchin, Anne O’Donnell-Luria, Steven M. Harrison, Sean V. Tavtigian, Marc S. Greenblatt, Leslie G. Biesecker, Predrag Radivojac, Steven E. Brenner, ClinGen Sequence Variant Interpretation Working Group

## Abstract

Recommendations from the American College of Medical Genetics and Genomics and the Association for Molecular Pathology (ACMG/AMP) for interpreting sequence variants specify the use of computational predictors as Supporting level of evidence for pathogenicity or benignity using criteria PP3 and BP4, respectively. However, score intervals defined by tool developers, and ACMG/AMP recommendations that require the consensus of multiple predictors, lack quantitative support. Previously, we described a probabilistic framework that quantified the strengths of evidence (Supporting, Moderate, Strong, Very Strong) within ACMG/AMP recommendations. We have extended this framework to computational predictors and introduce a new standard that converts a tool’s scores to PP3 and BP4 evidence strengths. Our approach is based on estimating the local positive predictive value and can calibrate any computational tool or other continuous-scale evidence on any variant type. We estimate thresholds (score intervals) corresponding to each strength of evidence for pathogenicity and benignity for thirteen missense variant interpretation tools, using carefully assembled independent data sets. Most tools achieved Supporting evidence level for both pathogenic and benign classification using newly established thresholds. Multiple tools reached score thresholds justifying Moderate and several reached Strong evidence levels. One tool reached Very Strong evidence level for benign classification on some variants. Based on these findings, we provide recommendations for evidence-based revisions of the PP3 and BP4 ACMG/AMP criteria using individual tools and future assessment of computational methods for clinical interpretation.

## INTRODUCTION

Genetic and genomic testing is now standard of care for identifying hereditary susceptibility to many conditions (e.g., cancer, metabolic conditions, intellectual and physical developmental disorders) as it can provide an etiologic diagnosis and indicate increased lifetime risk to manifest symptoms of a monogenic disease. However, testing can also identify variants of uncertain significance (VUS), many of which are amino acid substitutions.^1^ VUS are rapidly accumulating in variant databases and their classification represents a major challenge in clinical genetics.^1^

To help standardize the approach of clinical genetic/genomic testing laboratories, the American College of Medical Genetics and Genomics and the Association for Molecular Pathology (ACMG/AMP) published recommendations for evaluating the pathogenicity of variants in genes associated with monogenic disease.^2^ The ACMG/AMP recommendations (1) list qualitatively distinct lines of evidence (functional, genetic, population, computational, etc.), (2) indicate how each evidence type could be applied toward a Pathogenic or Benign classification, (3) stratify the strength of evidence as Supporting, Moderate, Strong, Very Strong, or Standalone for pathogenicity and benignity, and (4) provide rules for combining evidence types that defined the amount of evidence required to reach the classification categories.

Many computational (*in silico*) tools have been developed to predict if a variant will disrupt the function of a gene product.^3–5^ Because computational tools can be applied to many different types of genomic variation, these methods are attractive for application to variants observed in clinical or research testing, particularly in the absence of genetic or functional evidence. However, it is critical to recognize that while *in silico* predictors alone are not capable of classifying the pathogenicity of a variant, with adequate calibration and validation they can provide a useful contribution to the overall classification. The 2015 version of the ACMG/AMP computational classification rules stated that if “multiple lines of computational evidence” supported either pathogenic or benign classification, then they could be assigned the lowest level of evidence, “PP3-Supporting Pathogenic” or “BP4-Supporting Benign”. Supporting evidence must then be combined with substantial other lines of evidence to classify the variant as being pathogenic, benign, or of uncertain significance.

These rules have presented several challenges that can lead to either overstating or understating the strength of computational evidence.^3,6^ The ACMG/AMP recommendations required that two or more algorithms be used and that their outputs can be considered to be Supporting evidence only if the predictions from all tested algorithms agree. In practice, many methods overlap and thus do not offer independent assessment of pathogenicity.^4,6^ Accepted standards for concordance do not exist, and variability among laboratories has been observed.^7^ Finally, the design of several tools was motivated mainly by the discovery of novel variants and hypothesis generation for experimental follow-up, rather than clinical pathogenicity classification. As a result, the data sets used for validation and calibration of *in silico* predictors present potential sources of error for use of these tools in clinical settings, including the score thresholds required to apply evidence from any given predictor. Methods that have been tested on a few well-understood genes should be tested on larger data sets before they can be considered generalizable. If the variants from certain genes are overrepresented or if the same variants or different variants from the same protein occur both in the datasets used for training and for evaluation of these tools, the models may be overfitted and biased, and the effectiveness of the tool overestimated due to such circularities.^8^ Thus, the current ACMG/AMP rules can be applied by different labs in non-standardized ways that could lead to misestimation of the strength of *in silico* predictors for pathogenicity classification, encouraging inappropriate and/or inconsistent variant classification.

We have previously modeled the ACMG/AMP rules for combining evidence and showed how they fit a probabilistic framework.^9^ Under reasonable assumptions, we used a positive likelihood ratio of pathogenicity to quantify the strength of evidence that corresponds to a Supporting level of evidence and established that, within this framework, the strength of evidence required for Moderate, Strong, and Very Strong rose exponentially.^9^ This model gave us a basis for developing formal principles that can be used for validating and calibrating evidence for pathogenicity classification and potentially expanding the use of predictors beyond the Supporting evidence strength.^10,11^ This approach can also be used for *in silico* tools to establish proper weighting of computational evidence. With these, we can now study, using carefully curated data sets, how tools can be calibrated to support a given strength of evidence and can be used in the ACMG/AMP classification framework.

Here we propose a quantitative framework for establishing the level of contributory evidence in genomic testing that can be applied to any computational tool. We then focus on missense variation and systematically evaluate a set of widely used *in silico* tools on data sets validated to optimize accuracy and minimize circularity. We set out to determine score thresholds appropriate for a variant evaluated by the tool to reach various levels of evidence, potentially including levels beyond the original ACMG/AMP recommendation of Supporting. Our goal was to calibrate *in silico* tools, tested here for missense variants, so they could be used in a manner that is consistent across clinical diagnostic laboratories and properly weighted based on evidence. Finally, we discuss our findings and implications for an effective use of computational tools in the clinical interpretation of variants.

## MATERIALS AND METHODS

### *In silico* tools considered

Missense variant interpretation tools for this study were selected based on several factors, including their mention in the ACMG/AMP recommendations, their prevalence in current clinical workflows, their consistent performance in independent assessments such as the Critical Assessment of Genome Interpretation (CAGI), their contribution to methodological diversity, and ease to obtain prediction scores. This resulted in a set of 13 tools: BayesDel (without minor allele frequency),^12^ CADD,^13^ Evolutionary Action (EA),^14^ FATHMM,^15^ GERP++,^16^ MPC,^17^ MutPred2,^18^ PhyloP,^19^ PolyPhen-2 (HumVar model),^20^ PrimateAI,^21^ REVEL,^22^ SIFT,^23^ and VEST4.^24^ Of these, REVEL and BayesDel are meta-predictors and incorporate prediction scores from other tools, including some evaluated here. Except for BayesDel, CADD, EA, MutPred2, and PolyPhen-2, precomputed variant prediction scores for all tools were obtained from the database of human nonsynonymous SNPs and their functional predictions (dbNSFP).^25^ For the five tools mentioned above, prediction scores were generated for the variants considered in this study. All tools, except for FATHMM and SIFT, output scores such that higher scores are indicative of pathogenicity and lower scores are indicative of benignity. For consistency, the outputs of FATHMM and SIFT were transformed to be similar to the other tools and facilitate more consistent analyses. Note that we did not select any tool designed to evaluate splice variants.

#### Data sets

The main VCF file containing all variants from ClinVar^26^ was downloaded from its FTP site in December 2019. A series of steps were then undertaken to filter out variants that were not relevant to the analyses or could potentially bias them. Only missense variants with an allele frequency (AF) below 0.01 in the Genome Aggregation Database (gnomAD v2.1)^27^ and from genes with at least one pathogenic variant of any type in ClinVar were first retained. In this step, for each variant, the gnomAD exomes global AF was used. When this was unavailable, the gnomAD genomes global AF was used. Among these, all VUS, variants with a zero-star review status, i.e., without any detailed review information, and those with conflicting classifications were excluded. Next, variants that were present in the training sets of the different tools considered in this study were removed whenever available. Excluded variants came from the training sets of BayesDel, FATHMM, MPC, MutPred2, PolyPhen-2, REVEL, and VEST4. For meta-predictors such as BayesDel and REVEL that incorporate prediction scores from other tools, we also removed training variants for their constituent tools when available; e.g., FATHMM, MutPred,^28^ VEST3,^24^ and MutationTaster.^29^ The resulting data set consisted of 11,834 variants from 1,914 genes and is referred to as the *ClinVar 2019* set (Figure 1A).

**Figure 1.**
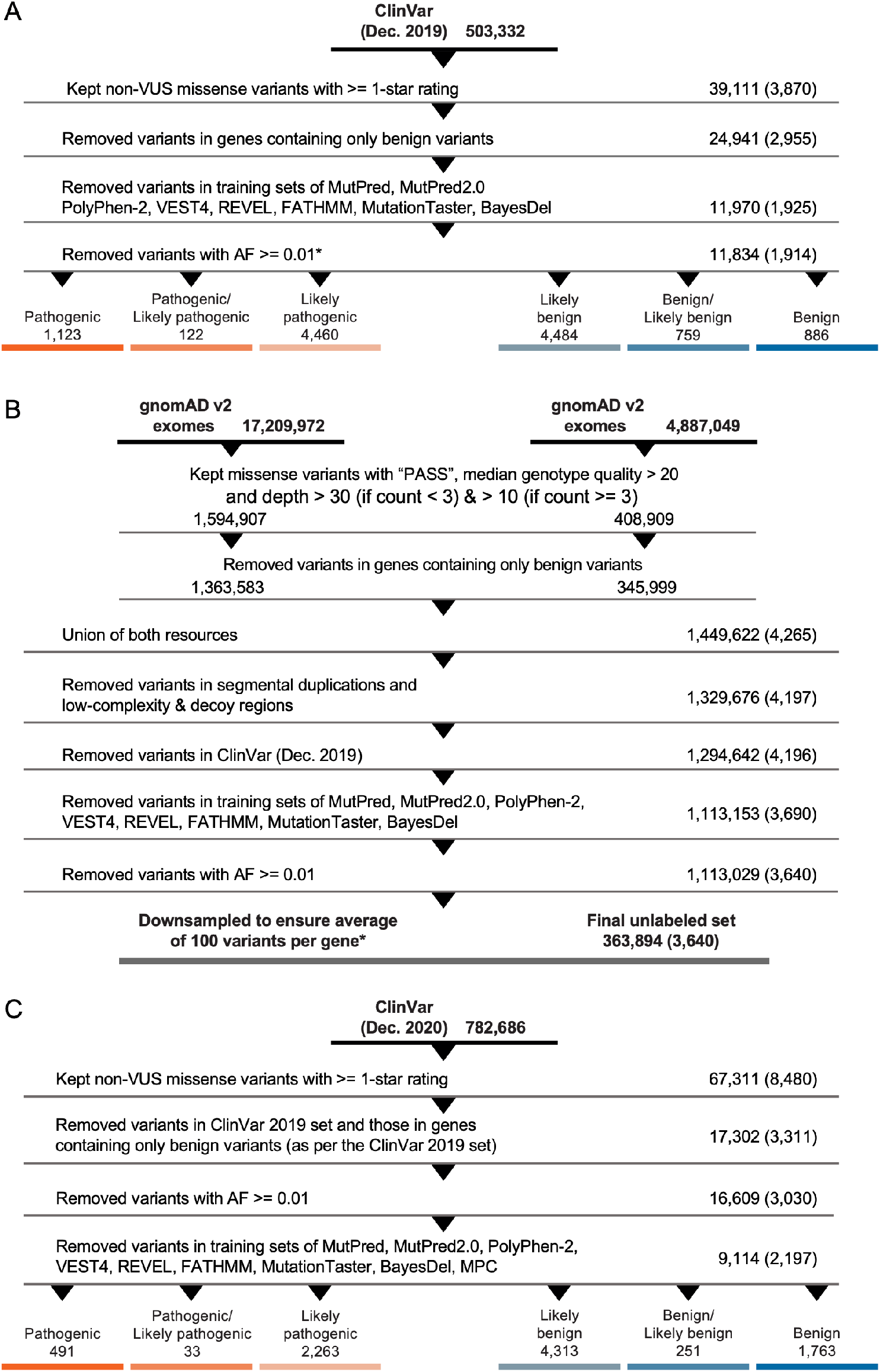
Data set preparation. Steps taken to prepare the three data sets in this study, extracted from ClinVar (A, C) and gnomAD (B). Numbers on the right side represent the numbers of variants remaining after each step and numbers in parentheses represent the numbers of genes remaining after each step. The data set resulting from (A) is referred to as the *ClinVar 2019* set, from (B) the *gnomAD* set, and from (C) the *ClinVar 2020* set. The asterisk refers to numbers after removing variants from the MPC training sets. This was done in a *post hoc* manner after all filtering and downsampling steps were carried out for the *ClinVar 2019* and *gnomAD* sets.

A second data set was created using VCF files from gnomAD. Both exome and genome data (v2.1.1) were obtained from the gnomAD downloads page, after which a series of filtering steps was undertaken. As before, only variants with AF < 0.01 were retained. For quality control, only those variants annotated as “PASS” in the “FILTER” column and with median genotype quality > 20 were retained. Additionally, all retained variants were required to have a median depth ≥ 30 if the allele count was < 3 (present in at most two individuals), or ≥ 10 if the allele count was at ≥ 3. As with the ClinVar set, all missense variants from genes without a single pathogenic variant of any type in ClinVar were removed. From this point on, data from both exomes and genomes were merged into a single data set of 1,449,622 variants by taking the union of the two resources. From these, variants in segmental duplications, low-complexity and decoy regions were removed. As before, variants present in the various predictors’ training sets were also removed. Finally, to ensure no overlap with the ClinVar set, gnomAD variants found in ClinVar (December 2019) were removed. The resulting data set consisted of 363,894 variants from 3,640 genes and is referred to as the *gnomAD* set (Figure 1B).

Finally, a test set consisting only of missense variants meeting the previously described criteria and present in ClinVar after December 2019 was created. The main VCF file containing all variants from ClinVar was downloaded from its FTP site in December 2020. All steps were identical to those undertaken for the *ClinVar 2019 set*, except that filtering against the tools’ training sets was undertaken at the end after all other filtering steps. The resulting data set was then cross-referenced against the *ClinVar 2019* set to obtain the final test set of 9,114 variants from 2,197 genes. This is referred to as the *ClinVar 2020* set (Figure 1C).

#### Estimation of the prevalence of pathogenic variants

Estimating the prevalence of pathogenic variants, or the prior probability of pathogenicity, requires selection of a reference set. We reasoned that rare variants in gnomAD among Mendelian disease genes constituted an appropriate set and then adopted a rigorous estimation approach using the AlphaMax algorithm.^30^ AlphaMax is a nonparametric maximum likelihood method that relied on a data set of ClinVar variants labeled as pathogenic or likely pathogenic and a gnomAD reference set of unlabeled variants. It typically maps high-dimensional input data into univariate output data, from which the priors are then estimated using kernel density estimation. To obtain a univariate transform, we trained a neural network-based method using the same input features and training steps as with MutPred2^18^ based on the result that neural networks can be good approximators of posterior distributions.^31^ This procedure yielded a prior probability of pathogenicity (prevalence in the data set) of 4.41%, higher than that estimated previously for an exome of a healthy individual^18^ using a similar procedure but lower than the prevalence of 10% assumed by Tavtigian et al.^9^ for clinical sequencing data.

#### Statistical framework

The ACMG/AMP recommendations suggested multiple levels of evidential strength to consider: Supporting, Moderate, Strong, and Very Strong for pathogenic and Supporting, Strong or Standalone for benign. Here, we have not considered the Standalone evidence for benignity further because variants with MAF > 0.05 were excluded from our data by default. We have added Moderate and Very Strong levels for benign in anticipation of future need for such strengths although our work here is applicable to both symmetric and asymmetric notions of strength for pathogenicity and benignity. In accordance with a recent proposal,^32^ we included an additional category of Indeterminate for variants not reaching evidential strength of Supporting for either pathogenicity or benignity. Thus, the indeterminate variants for a given tool would not add evidence strength to variant classification. Note that predictors differed in the number and identity of indeterminate variants (see Results section). To reduce subjectivity and define strengths of evidence in a quantitative framework, these were mapped to odds ratios of pathogenicity or positive likelihood ratios,^33^ as in Tavtigian et al.,^9^ so that the posterior probability of combined evidence listed in the ACMG/AMP recommendations for a likely pathogenic variant was at least 0.9 and less than 0.99, and for a pathogenic variant at least 0.99. Similarly, we required identical values for the posterior probability of benignity, that corresponded to the thresholds of 0.1 and 0.001 for the posterior probability of pathogenicity. However, as noted above, the estimated prevalence of pathogenic variants in our *gnomAD* reference set was 4.41%. vs. the clinical experience-based 10% value primarily considered in Tavtigian et al.^9^

We started by connecting the posterior odds of pathogenicity given the evidence and the positive likelihood ratio (LR^+^) using the following expression

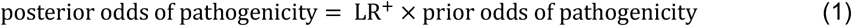

where, for a variant ν, and on a particular reference data distribution

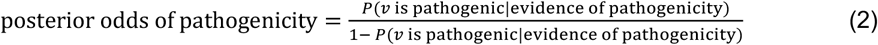

and

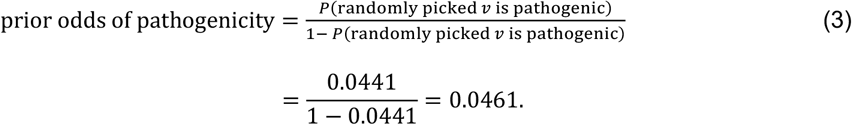

In Equations 2 and 3, *P*(ν is pathogenic|evidence of pathogenicity) is the posterior probability of pathogenicity and *P*(randomly picked ν is pathogenic) is the prior probability of pathogenicity on a reference set. When considering computational methods, the “evidence of pathogenicity” corresponds to a discretized prediction that the variant is pathogenic.

It can be shown^33^ that LR^+^ is independent of the class prior and can be alternatively expressed as

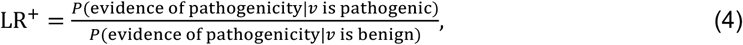

which has a straightforward interpretation for binary classification models, because *P*(evidence of p*a*thogenicity|ν is p*a*thogenic) is the true positive rate and *P*(evidence of p*a*thogenicity|ν is benign) is the false positive rate.

To incorporate the combining nature of multiple lines of evidence and model the ACMG/AMP rules with few parameters, Tavtigian et al. sought to express LR^+^ in an exponential form

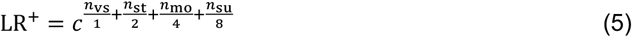

where *n*_vs_, *n*_st_, *n*_mo_, and *n*_su_ are the number of Very Strong, Strong, Moderate, and Supporting lines of evidence, respectively. The value of *c* can be determined either computationally or manually so that the ACMG/AMP rules are generally satisfied in that the posterior probability reaches the values of 0.9 and 0.99 for likely pathogenic and pathogenic classifications, respectively. More importantly, one can readily verify from Equation 5 that a single line of Very Strong, Strong, Moderate, and Supporting evidence must reach 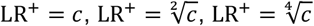, and 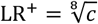, respectively. Given the prior probability of pathogenicity of 0.0441, we obtained *c* = 1,124, from which we further obtained the LR^+^ values of single lines of evidence (Table 1).

**Table 1.** Posterior probability and positive likelihood ratio values (reduced to four significant digits) that define the varying strengths of evidence in this study for the PP3 and BP4 criteria.

| Criterion | Evidence Strength | Posterior probability of pathogenicity (for PP3) and benignity (for BP4) | Positive likelihood ratio | Reciprocal of positive likelihood ratio |
| --- | --- | --- | --- | --- |
| PP3 | Very Strong | 0.9811 | 1,124:1 | 0.0009:1 |
|  | Strong | 0.6073 | 33.53:1 | 0.0298:1 |
|  | Moderate | 0.2108 | 5.790:1 | 0.1727:1 |
|  | Supporting | 0.0999 | 2.406:1 | 0.4156:1 |
| BP4 | Supporting | 0.9812 | 0.4156:1 | 2.406:1 |
|  | Moderate | 0.9921 | 0.1727:1 | 5.790:1 |
|  | Strong | 0.9986 | 0.0298:1 | 33.53:1 |
|  | Very Strong | 1.0000 | 0.0009:1 | 1,124:1 |

Most computational tools, however, do not discretize their predictions and instead only provide a raw score *s* ∈ ℝ for a given variant ν, thus leaving it up to the variant analyst to interpret the score and define an appropriate threshold. This suggested that in such cases we needed to use a continuous score *s* as evidence of pathogenicity for which Equation 4 breaks down. To address this, we defined a local positive likelihood ratio lr^+^, an equivalent of LR^+^ that could be used with continuous evidence, as a density ratio between score distributions on pathogenic and benign variants; that is,

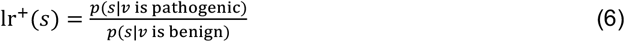

and sought to estimate it from prediction data on a set of variants for each considered tool. From here, we computed the local posterior probability values as

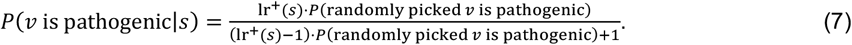

Equation 7 can be derived from Equations 1, 2 and 3 with lr^+^ in place of LR^+^. To contrast LR^+^ with lr^+^, we will refer to LR^+^ as the global likelihood ratio. Note further that the posterior probability and the likelihood ratio can be expressed as functions of each other and the prior probability of pathogenicity. In this study, since the same prior probability of pathogenicity (0.0441) is used for all analyses, local posterior probabilities and local likelihood ratios are considered equivalent and are used interchangeably. The density ratio and the posterior can be practically estimated using a narrow sliding window around each score value *s* (Figure 2). This approach is sound and the local posterior corresponds to the local positive predictive value for a computational tool, an equivalent of the local false discovery rate.^34,35^

**Figure 2.**
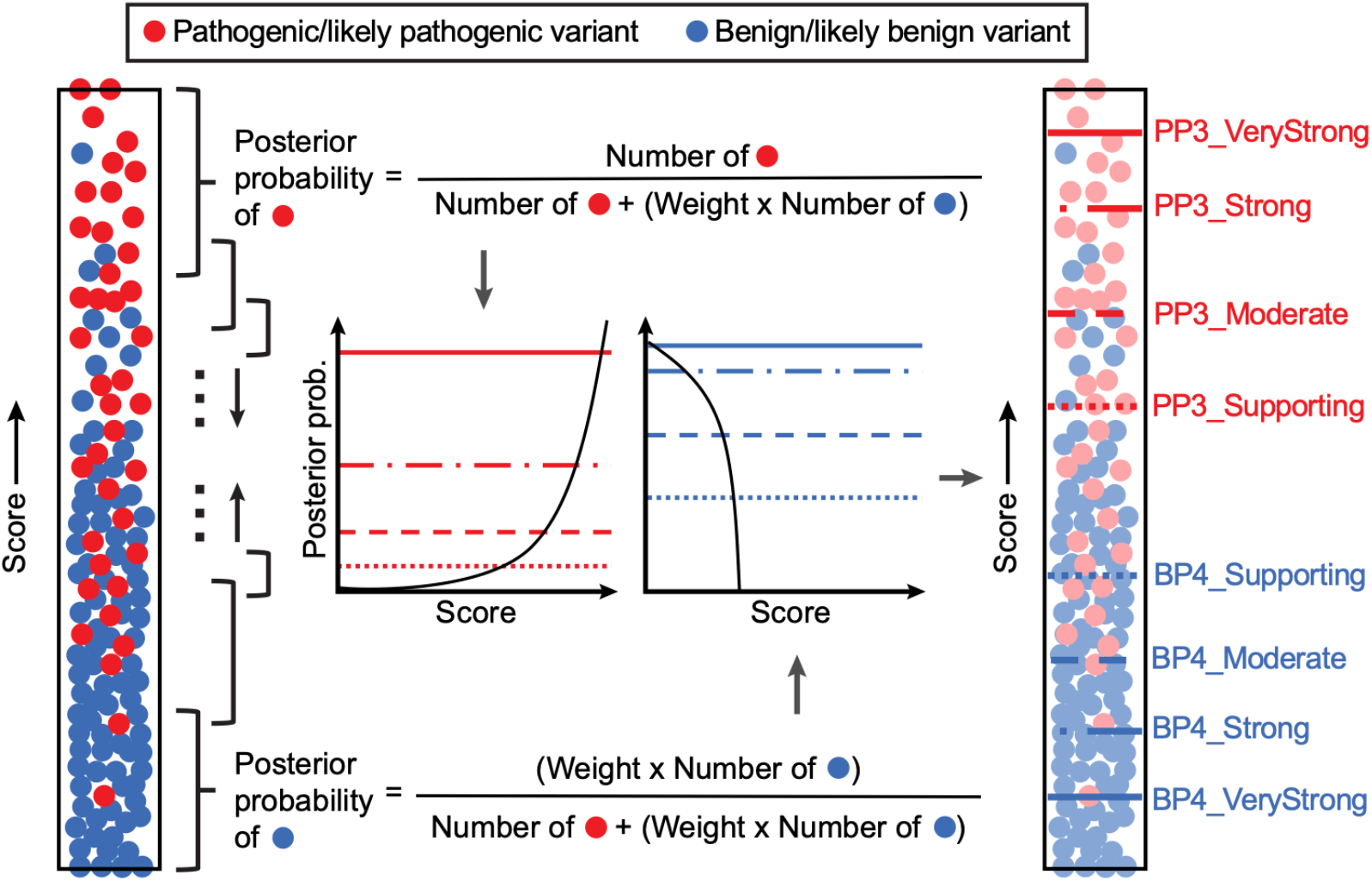
Conceptual representation of the estimation of intervals for evidential support. An example *in silico* tool that is supposed to assign higher scores to pathogenic variants is shown. Each filled circle represents a variant, either pathogenic/likely pathogenic (red) or benign/likely benign (blue) as recorded in the *ClinVar 2019* set. All unique scores were first sorted and each score was then set as the center of the sliding window or the local interval (black-colored braces), within which posterior probabilities were calculated. Here, to ensure that a sufficient number of variants were included in each local interval, *ϵ* was adaptively selected to be the smallest value so that the interval [*s* − *ϵ, s* + *ϵ*]around a prediction score *s* incorporated at least 100 pathogenic and benign variants (combined) from the *ClinVar 2019* set and at least 3% of rare variants from the *gnomAD* set with predictions in the given local interval, separately for each method (technically, *ϵ* is a function of score *s* for each predictor). These numbers were proportionally scaled at the ends of the score range. The estimated posterior probabilities were then plotted against the output scores. Using posterior probability thresholds defined in Table 1, score thresholds were subsequently obtained for pathogenicity (PP3) and benignity (BP4) for each method. Here, the number of benign variants was weighted to calibrate methods according to the prior probability of pathogenicity. The weight was calculated by dividing the ratio of pathogenic and benign variant counts in the full data set by the prior odds of pathogenicity; see Eq. 3. The pathogenic and benign counts (and this weight) slightly varied for each method because scores were not available for all variants in the data set for some tools. In this study, the estimated prior probability of pathogenicity (0.0441) was used to account for the enrichment of pathogenic/likely pathogenic variants in ClinVar. The estimated prior probability of benignity was assumed to be 1 – 0.0441 = 0.9559.

#### Computational objective

The objective of our approach was to develop a framework that discretizes the prediction range for any given computational tool into a set of nine intervals, corresponding to the various levels of evidential strength: Supporting, Moderate, Strong, and Very Strong for both benign and pathogenic, and Indeterminate (for the scores not satisfying any of the levels of desired evidential strength). To do so, we considered an *in silico* tool, or a scoring function, that outputs a pathogenicity score *s* ∈ ℝ for a variant ν where the higher scores were designed to suggest stronger evidence for pathogenicity than the lower scores. We searched for a set of score thresholds 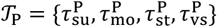 such that a prediction score 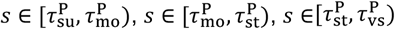, and 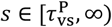 could be considered Supporting, Moderate, Strong, and Very Strong evidence for pathogenicity, respectively. We will refer to 𝒯_p_ as the pathogenicity threshold set and for convenience, we will refer to the above-mentioned contiguous prediction intervals as 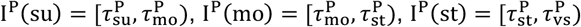, and 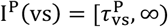. Equivalently, the evidence supporting benignity upon seeing a prediction output *s* requires us to determine the benignity threshold set 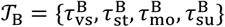, with the interpretation that 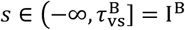 is the Very Strong evidence level for benignity, etc. Variants with predictions 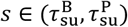 were considered to lie in the indeterminate region, thus neither supporting pathogenicity nor benignity for such variants. Without loss of generality, we assumed that each predictor reached all four levels of evidential support for pathogenicity and benignity; however, this was not the case in practice as the evidence levels achieved by different predictors depended on the characteristics of each tool’s score distributions.

#### Estimating intervals for levels of evidential support

We then turned to determining the threshold sets 𝒯_p_ and 𝒯_B_ for each model to establish a set of up to nine intervals (four for pathogenicity, four for benignity, and the indeterminate region, see Table 1). We focused on the set 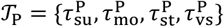 first, which defined the contiguous intervals I^p^(evidence level), where evidence level ∈ {su, mo, st, vs}. To define the threshold set 𝒯_p_ and, ultimately, the pathogenic intervals I^p^(evidence level), we defined the threshold for the Supporting level of evidence as

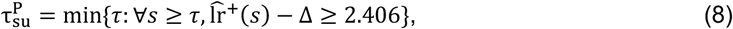

where 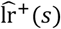 is estimated lr^+^(*s*) using an *ϵ* neighborhood around *s*; that is, all prediction scores *s* ∈ [*s* − *ϵ, s* + *ϵ*] are considered pathogenic and used to compute lr^+^(*s*) and the local positive predictive value. The parameter Δ is a nonnegative margin of error selected so that 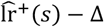 is the value of the one-sided 95% confidence bound of the estimated lr^+^(*s*), and was determined via 10,000 bootstrapping iterations. In other words, the threshold for the Supporting level of evidence was the smallest value *τ* in the prediction range such that for all scores *s* greater than or equal to *τ*, the lower error bound of 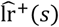 was greater than or equal to 2.406. The remaining thresholds from 𝒯_p_ are determined in the same manner, and the procedure is then repeated for 𝒯_B_ using

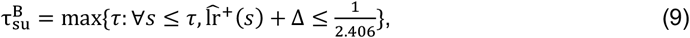

and so on. The parameter Δ was incorporated to lead to a more stringent threshold selection.

#### Validating the local approach for interval estimation

Variants in the *ClinVar 2020* set and the *gnomAD* set were used to determine if our local approach for the estimation of strength-based intervals was robust. For each tool, the thresholds selected using the above procedure were applied to assign each variant into an interval of evidential strength. Within each interval, two measures were computed: (1) an interval-based likelihood ratio was calculated to verify if our estimated intervals did indeed provide evidence for pathogenicity/benignity with the expected strength on variants not seen by our estimation procedure (*ClinVar 2020* set), and (2) the fraction of variants in the *gnomAD set* that fell into each interval was calculated to assess overprediction of pathogenic variants, particularly at higher evidential strengths.

The interval-based likelihood ratio is simply the global likelihood ratio calculated over a given score interval:

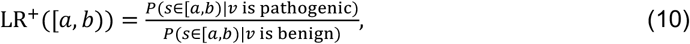

where *a* and *b* are the lower and upper bound of the score interval, respectively, and *s* and ν are defined as above. In this case, rather than considering a variant’s score directly or a binarized version of it as the evidence for pathogenicity/benignity, the categorization of the variant into one of the strength categories was used as evidence. We expected 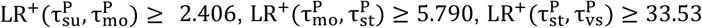, and 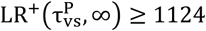. In other words, the interval-based likelihood ratio had to be greater than or equal to that obtained using the local likelihood ratio approach. This should have held for benignity intervals as well. Here, each interval [*a, b*) was instantiated from the sets of optimal thresholds obtained using the local likelihood ratio approach, 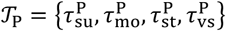 and 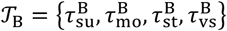. The interval-based likelihood ratio was then operationalized as the ratio of the true positive rate to the false positive rate within the interval using the *ClinVar 2020* set. Depending on whether the interval was for pathogenicity or benignity, the true positive rate was either the fraction of pathogenic/likely pathogenic variants falling within the interval or the fraction of benign/likely benign variants. Similarly, the false positive rate was the fraction of benign/likely benign variants and the fraction of pathogenic/likely pathogenic variants within the interval, respectively.

## RESULTS

### *In silico* tools yield levels of evidence beyond Supporting

Our local posterior probability-based approach allowed for the systematic identification of thresholds corresponding to different strengths of evidence for a given tool. To this end, using the *ClinVar 2019* set, we applied this approach on thirteen tools: BayesDel,^12^ CADD,^13^ EA,^14^ FATHMM,^15^ GERP++,^16^ MPC,^17^ MutPred2,^18^ PhyloP,^19^ PolyPhen-2,^20^ PrimateAI,^21^ REVEL,^22^ SIFT,^23^ and VEST4.^24^ We first obtained scores from these tools for variants in this data set and then calculated local posterior probabilities for each unique score, as shown in Figure 2. Scores that satisfied the posterior probability thresholds presented in Table 1 were then deemed to provide the corresponding strengths of evidence (Table 2).

**Table 2.** Estimated threshold ranges for all tools in this study corresponding to the four pathogenic and four benign intervals. A “–” implies that the given tool did not meet the posterior probability (likelihood ratio) threshold. See Supplemental Table S1 for comprehensive results that include point estimates and one-sided confidence intervals. Intervals follow standard mathematical notation in which “(“and “)” indicate exclusion of the end value and “[“and “]” indicate inclusion of the end value.

| Method | Pathogenic (PP3) |  |  |  | Benign (BP4) |  |  |  |
| --- | --- | --- | --- | --- | --- | --- | --- | --- |
|  | Very Strong | Strong | Moderate | Supporting | Supporting | Moderate | Strong | Very Strong |
| BayesDel | - | ≥ 0.50 | [0.27, 0.50) | [0.13, 0.27) | (-0.36, -0.18] | ≤ -0.36 | - | - |
| CADD | - | - | ≥ 28.1 | [25.3, 28.1) | (17.3, 22.7] | (0.15, 17.3] | ≤ 0.15 | - |
| EA | - | - | ≥ 0.821 | [0.685, 0.821) | (0.069, 0.262] | ≤ 0.069 | - | - |
| FATHMM | - | - | ≤ -5.04 | [-5.04, -4.14) | (3.32, 4.69] | ≥ 4.69 | - | - |
| GERP++ | - | - | - | - | (-4.54, 2.70] | ≤ -4.54 | - | - |
| MPC | - | - | ≥ 1.828 | [1.360, 1.828) | - | - | - | - |
| MutPred2 | - | ≥ 0.932 | [0.829, 0.932) | [0.737, 0.829) | (0.197, 0.391] | (0.010, 0.197] | ≤ 0.010 | - |
| PhyloP | - | - | ≥ 9.741 | [7.367, 9.741) | (0.021, 1.879] | ≤ 0.021 | - | - |
| PolyPhen2 | - | - | ≥ 0.999 | [0.978, 0.999) | (0.009, 0.113] | ≤ 0.009 | - | - |
| PrimateAI | - | - | ≥ 0.867 | [0.790, 0.867) | (0.362, 0.483] | ≤ 0.362 | - | - |
| REVEL | - | ≥ 0.932 | [0.773, 0.932) | [0.644, 0.773) | (0.183, 0.290] | (0.016, 0.183] | (0.003, 0.016] | ≤ 0.003 |
| SIFT | - | - | ≤ 0.000 | [0.000, 0.001) | (0.080, 0.327] | ≥ 0.327 | - | - |
| VEST4 | - | ≥ 0.965 | [0.861, 0.965) | [0.764, 0.861) | (0.302, 0.449] | ≤ 0.302 | - | - |

We were able to identify thresholds for Supporting and Moderate levels of evidence for pathogenicity (PP3) and benignity (BP4) for all tools, except for GERP++, which did not yield Supporting evidence for PP3, and MPC, which did not yield Supporting evidence for BP4. Interestingly, the local posterior probability curves showed that, at appropriate thresholds, several tools could provide Strong evidence for pathogenicity (BayesDel, VEST4), benignity (CADD) or both (MutPred2, REVEL), as shown in Figure 3, Table 2, and Supplemental Table S1.

**Figure 3.**
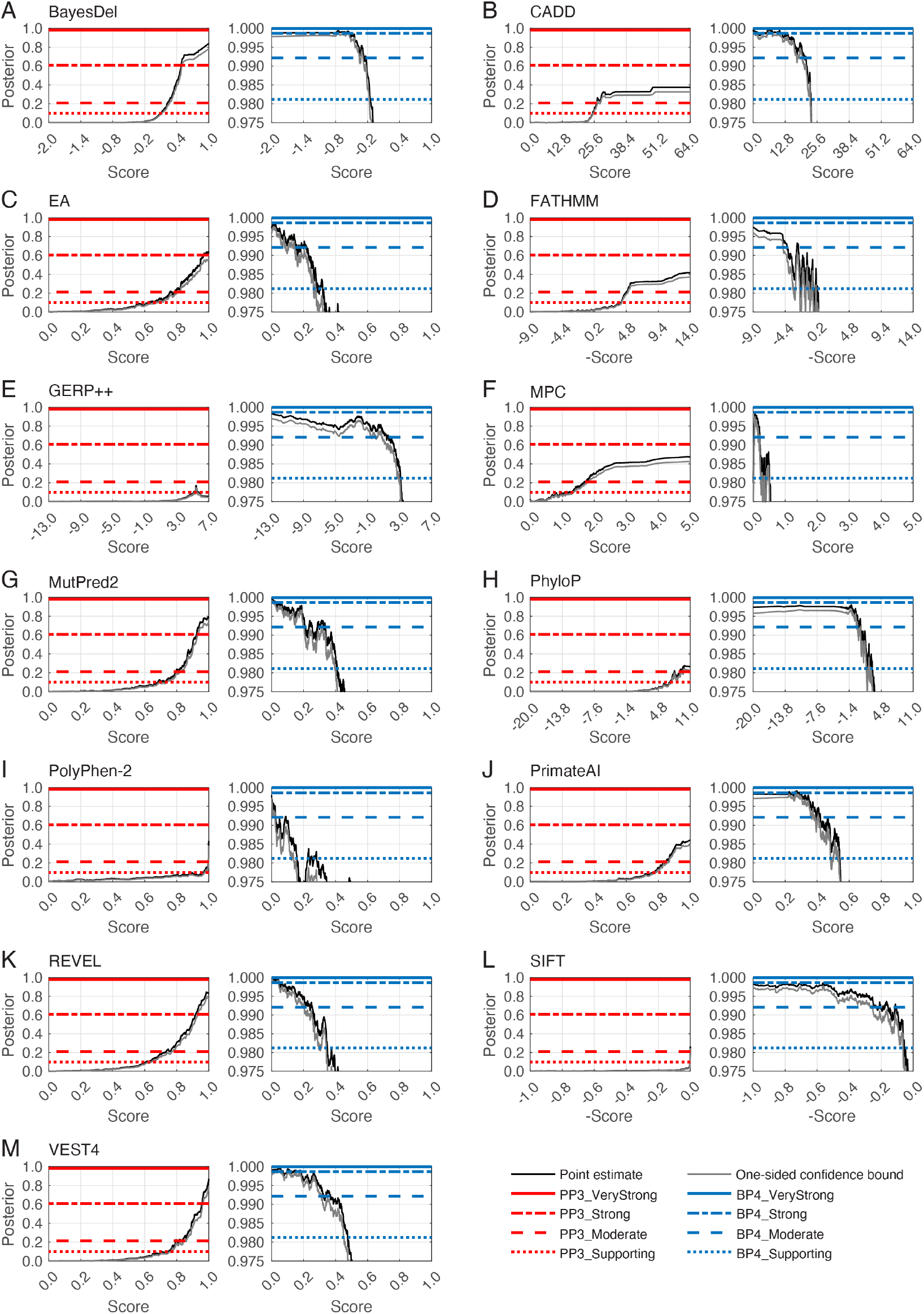
Local posterior probability curves for (A) BayesDel, (B) CADD, (C) Evolutionary Action (EA), (D) FATHMM, (E) GERP++, (F) MPC, (G) MutPred2, (H) PhyloP, (I) PolyPhen-2, (J) PrimateAI, (K) REVEL, (L) SIFT, and (M) VEST4. For each panel, there are two curves: the curve on the left is for pathogenicity (red horizontal lines) and the curve on the right is for benignity (blue horizontal lines). The horizontal lines represent the posterior probability thresholds for Supporting, Moderate, Strong, and Very strong evidence. The black curves represent the posterior probability estimated from the *ClinVar 2019* set. The grey curves represent one-sided 95% confidence intervals calculated from 10,000 bootstrap samples of this data set (in the direction of more stringent thresholds). The points at which the grey curves intersect the horizontal lines represent the thresholds for the relevant intervals.

#### Validation of estimated score intervals on an independent data set

Two approaches were adopted to assess the correctness of our local posterior probability strategy. First, we verified if the intervals that we estimated corresponded to similar lr^+^ values on an independent data set as did those on the estimation data set. We reasoned that the likelihood ratios in each interval should equal or exceed those estimated using the sliding window algorithm. To this end, we classified variants in the *ClinVar 2020* set into each of the four intervals for pathogenicity and benignity, and calculated likelihood ratios within each interval. We found that all methods that reached the Strong level of evidence for pathogenicity exceeded the likelihood ratio value in Table 1 (Figure 4A). In addition, all tools that met the threshold for Moderate levels of evidence for pathogenicity resulted in likelihood ratios exceeding the corresponding likelihood ratio threshold required, except for PhyloP which had a marginally lower likelihood ratio. Similarly, all tools that met the Supporting level of evidence on the *ClinVar 2019* data exceeded the likelihood ratio threshold for the Supporting evidence interval of 2.41 on the *ClinVar 2020* data. For benignity, a tool must have had an interval-based likelihood ratio lower than those in Table 1 for the relevant evidential strength, a criterion met by all tools.

**Figure 4.**
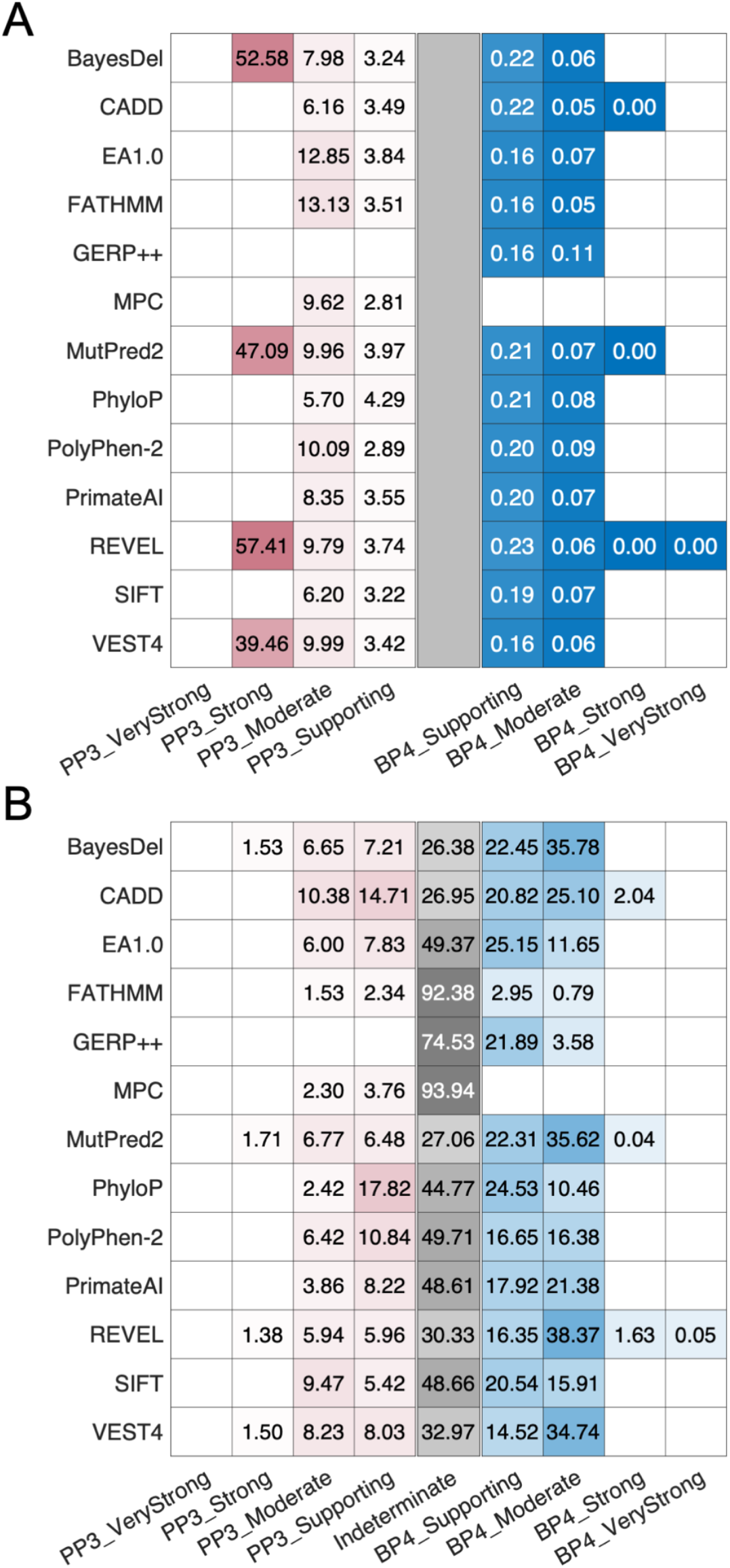
Evaluation of the robustness of our approach and estimated score intervals. (A) The likelihood ratios within each interval on the independent *ClinVar 2020* set. (B) The percentage of variants predicted to be within the interval in the *gnomAD* set. Red and blue distinguish between the evidential strength intervals for pathogenicity and benignity, respectively, with the indeterminate interval colored grey. The color gradient corresponds to the value in the cells, regardless of color. In (A), darker colors indicate higher values for pathogenicity and lower values for benignity (because these are positive likelihood ratios), and in (B), darker colors indicate higher proportions. A grey rectangle is introduced at the center of (A) for comparability across the two panels. White cells without values indicate that the tool did not yield thresholds corresponding to the relevant intervals. The indeterminate interval in (B) also included variants without any scores. For each tool, the fraction of variants with missing predictions is reported in Supplemental Table S2. When interpreting these findings, the totality of the results in (A) and (B) must be considered to account for the effects of binning of continuous scores into discrete intervals. For example, although a tool such as CADD provides most predictions classified to be Supporting and Moderate for PP3 (B), it does so with lower accuracy (A), measured by the smaller number of true positive predictions for the same number of false positive ones, than a tool such as REVEL. Due to the effects of binning, many of the true positive predictions for REVEL are in its Strong evidence category, further obscuring interpretation. Thus, the results in Table 2 and this figure must be considered with utmost care for any use outside our recommendations; see below.

Second, we verified that these intervals were stringent in their assignment of Strong levels of pathogenicity to variants in the *gnomAD* set. We reasoned that the prior probability of a pathogenic variant in a population such as that of the *gnomAD* set is low. Therefore, the Strong pathogenic score interval should ideally classify only a small fraction of variants as Strong. We found that for tools that reached the Strong level of evidence for pathogenicity, the fraction of gnomAD variants with Strong levels of evidence ranged from 1.4 to 1.7% (Figure 4B). These fractions were smaller than the experience-based prior probability assumed by Tavtigian et al. (10%) and the prior probability that was estimated in this study using the *gnomAD* set (4.41%). Furthermore, when considering the Strong and Moderate intervals together, all tools except CADD yielded a smaller fraction of variants than 10%. Interestingly, for benignity, 6 out of 13 tools assigned a majority of variants to Moderate rather than Strong or Supporting evidence. Most tools output a score in the Indeterminate range (or did not have scores) for 26-50% of variants and would not provide any evidence strength for classification of these variants. FATHMM, GERP++ and MPC were outliers, resulting in more than half of variants in the Indeterminate range. Taken together, these results suggest that our local posterior strategy generally yields robust intervals that are unlikely to overestimate the evidential strength that they provide.

#### Investigation of developer-recommended thresholds and simple consensus approaches

We investigated whether *in silico* tools that are frequently used in clinical variant interpretation and their default score thresholds, met the quantitative definition of Supporting evidence as described in Table 1. We focused on the local posterior probability curves and score intervals estimated above using the *ClinVar 2019* data set for three tools and their combinations: SIFT, PolyPhen-2 and CADD. We then checked the local likelihood ratio values at the developer-recommended thresholds against those in Table 1 to assess the strength of evidence obtained at these thresholds.

At the developer-recommended thresholds, SIFT, PolyPhen-2, and CADD did not meet the local likelihood ratio thresholds for Supporting evidence for PP3 (Table 3). Interestingly, the developer-recommended threshold of 20.0 for CADD was in the score interval corresponding to Moderate evidence for BP4 (Figure 3; Table 2), suggesting an inappropriate use of this threshold as evidence for pathogenicity. Furthermore, all tools classified a substantial fraction of variants in the *gnomAD* set as damaging (50.4% by SIFT, 29.3% by PolyPhen-2 and 65.1% by CADD) at developer-recommended thresholds. These fractions were considerably larger than 4.41%, our estimate of the prior probability of pathogenicity (prevalence of pathogenic variants) in the *gnomAD* set, suggesting a high false positive rate with respect to PP3, in agreement with previous studies.^3^

**Table 3.** Assessment of the strength of evidence for PP3 through local likelihood ratios provided by SIFT, PolyPhen-2, and CADD using default developer-recommended thresholds. Numbers in parentheses indicate lower and upper (95%) confidence intervals as calculated by the bootstrapping method. “Evidence strength” was determined on the basis of the point estimate of the positive likelihood ratio. The Pearson correlation between outputs of SIFT and PolyPhen-2 on our *gnomAD* set was 0.47, the correlation between SIFT and CADD was 0.49, and the correlation between PolyPhen-2 and CADD was 0.67. Correlation between other tools is provided in Supplemental Figure S1.

| Method | Pathogenicity score threshold | $lr^+$ | Evidence strength | Fraction of predicted pathogenic variants in the *gnomAD* set |
| --- | --- | --- | --- | --- |
| SIFT | 0.050 | 0.048<br>(0.025, 0.075) | Not met | 0.504 |
| PolyPhen-2 | 0.902 | 1.993<br>(1.560, 2.492) | Not met | 0.293 |
| CADD | 20.000 | 0.157<br>(0.107, 0.215) | Not met | 0.651 |

We also implemented a simple consensus-based predictor using these three tools, emulating an approach typically adopted in clinical laboratories. We considered all possible pairs of tools and all three tools together by estimating lr^+^(***s***), where ***s*** is a two-dimensional score ***s*** = (s_1_, s_2_) or three-dimensional score ***s*** = (*s*_1_, *s*_2_, *s*_3_). We found that no combination of the three tools met the Supporting level of evidence PP3 using developer-recommended thresholds, although PolyPhen-2 individually, with score 0.902, was close to the desired lr^+^(*s*) threshold of 2.41. It is worth mentioning that this does not mean that there was no combination of scores for which these tools did not reach the Supporting evidence level, but rather that the appropriate evidential support was not met when all tools predicted scores at, or slightly better than, the developer-recommended minimum scores (or maximum, for SIFT). To the best of our knowledge, separate developer-recommended thresholds for predictions of benignity for these tools do not exist, and therefore, could not be evaluated.

## DISCUSSION

Our results provide the basis for refining how computational tools can be used to provide evidence for or against pathogenicity of variants using the Bayesian adaptation of the ACMG/AMP framework. The thresholds that we calculated show that these tools can provide stronger than Supporting evidence and that computational tools varied in their ability to reach these levels of evidence.

### Recommendations for updates to PP3 and BP4 criteria

For missense variants, to determine evidence for codes PP3 and BP4, we recommend that, for most situations, clinical laboratories use a single tool, genome-wide, that can reach the Strong level of evidence for pathogenicity and Moderate for benignity (BayesDel, MutPred2, REVEL, VEST4 among the tools evaluated here). This recommendation maximizes the strength of evidence that can be applied while minimizing the number of false positive predictions in the Supporting and Moderate categories. However, any method described here is valid to use for *in silico* evidence for PP3 and BP4 at the thresholds described and their relevant strengths. The choice of which tool to use for PP3/BP4 missense evidence must always be made before seeing prediction results and preferably other lines of evidence, to avoid biases such as those from multiple trials.

In situations where Variant Curation Expert Panels (VCEPs), clinical laboratories or research groups have developed gene-specific guidance, such as for the *RYR1* gene in malignant hyperthermia,^11^ laboratories could select the recommended alternative single tool and thresholds for these variants, instead of their standard tool otherwise used in genome-wide application. The importance of selecting a single tool to use for PP3/BP4 missense evidence is to avoid biases that could be introduced by, for example, scanning multiple tools for the strongest evidence for a given variant. We encourage calibration of tools for specific genes and regions using the methods described herein, with attention to the distinguishing features described below.

We have not evaluated the use of combining missense impact prediction methods for PP3/BP4 with methods that predict other mechanisms of genetic variant impact (e.g., splicing, expression) that could also be reported as PP3/BP4. However, computational methods that predict mechanistic consequences of the missense event (e.g., protein stability) should not be combined with other missense impact predictors, such as those evaluated here. Additionally, we have introduced caveats about combining rules for the moderate and strong PP3/BP4 with other evidence codes, as described below. The proposed recommendations for the use of computational tools, contrasted with those from the 2015 ACMG/AMP recommendations, are summarized in Table 4.

**Table 4.** Summary of recommendations for updates to PP3 and BP4 criteria in the ACMG/AMP recommendations and comparison with the 2015 ACMG/AMP recommendations.

| **Guidance** | **2015 ACMG/AMP recommendations** | **Our recommendation** |
| --- | --- | --- |
| Evidential strength of *in silico* tools | Supporting. | PP3_Strong, PP3_Moderate, PP3_Supporting, Indeterminate, BP4_Supporting, BP4_Moderate, and BP4_Strong. |
| Tools to be used | No specific recommendation, although a list of tools provided with the caveat that they have not been validated. | Tools that reach at least Strong evidence for pathogenicity and Moderate for benignity. It is valid to use any tool at the indicated evidential strength level as determined by our approach, so long as it is chosen before seeing its scores and preferably any other evidence. |
| Score thresholds for tools | No explicit recommendations but often defaults to developer-recommended thresholds. | Specific score thresholds calibrated to different evidential strengths. |
| Number of tools to be used | Multiple lines of evidence, suggesting multiple tools (no number provided). | Combinations of tools was not systematically assessed but results suggest the use of a single tool rather than an uncalibrated consensus of multiple tools. |
| Decision-making | If all agree, evidence can be counted as Supporting. Otherwise, evidence from tools cannot be used. | Evidence can be counted at the appropriate strength depending on the tool and score threshold chosen. The maximal strength of a tool must be committed to, i.e., if a tool provides at most PP3_Moderate strength and a variant satisfies the corresponding score threshold, it can only be counted as Moderate evidence. |
| Combing rules with other evidence codes | PP3 and BP4 can be applied for supporting evidence without consideration of other codes used. | Limit the combined evidence from PM1 and PP3 to Strong. |

We have invoked a level of evidence strength that was not included in the original ACMG/AMP recommendations (Benign Moderate, reflected by BP4_Moderate). This evidence level, while straightforward to derive from the data we present here, is not compatible with Table 5 “Rules for combining criteria to classify sequence variants” in the ACMG/AMP recommendations.^2^ There are several pragmatic, interim solutions for this issue. The first is that if a variant is shown to have BP4_Moderate evidence, Table 5 can be adapted by allowing that Likely Benign combining rule i, which is “1 Strong (BS1–BS4) and 1 supporting (BP1–BP7)”, be invoked if there is a strong benign criterion met (BS1-BS4) and use of an *in silico* tool generates BP4_Moderate. For the second Likely Benign combining rule (ii), it could be invoked if the only evidence generated is BP4_Moderate or if BP4_Moderate evidence is present along with an additional Benign supporting evidence type (BP1-BP3, BP5, or BP7). Other combinations of BP4_Moderate evidence are not germane to those combining rules. A second approach would be to use the Tavtigian et al. framework^9^, in either its initial iteration (see Supplemental Methods for suggested modifications), or in its simplified, points-based iteration,^32^ in which BP4_Moderate evidence would count as −2 points toward variant classification.

### Alternative strategies for interval definition

We selected the local approach as our main strategy owing to its theoretical foundation and simplicity compared with other statistical methods. We investigated two alternative strategies to define intervals corresponding to the relevant evidential support. The first alternative strategy calculated the global positive likelihood ratio over the entire data set above (for PP3) or below (for BP4) a certain score; see Supplemental Methods. Score thresholds were then defined in the same manner as those in the local approach, i.e., by selecting the desired likelihood ratio levels from Table 1, yielding a global interpretation for evidential strength. However, this strategy had two critical flaws: (1) each desired likelihood ratio level was satisfied only on *average* within any interval, likely misclassifying the evidence strengths for variants at the extremes of the range; and (2) unlike the local approach, it did not guarantee that the threshold for PP3_Supporting was always higher than BP4_Supporting, resulting in difficulties of reconciling the threshold sets obtained in this way.

The second alternative strategy for selecting threshold sets sought to estimate all thresholds simultaneously by optimizing the interval-based likelihood ratios in Equation 10 directly, such that each interval resulted in a positive likelihood ratio value equal to or greater than the corresponding one in Table 1. This approach, however, had three significant flaws: (1) it may not have yielded a unique solution and further required an optimization algorithm to select thresholds; (2) the threshold set was determined jointly and thus each resulting interval of evidential support depended on the number of levels for evidential strength prescribed by the ACMG/AMP recommendations, which could change in the future, altering the results; and (3) as above, the likelihood ratios were still determined on average within intervals.

Therefore, we selected the local approach, which is also the most stringent approach to determining the thresholds (with respect to avoiding overestimation of evidence) as the appropriate evidence level holds for any score in the determined evidence interval.

### Implications for the assessment of additional clinical variant interpretation tools, including those developed in the future

Several studies have attempted to benchmark methods to identify tools and tool-threshold combinations that are the most accurate and appropriate for clinically relevant variants.^3,36–39^ Key distinguishing aspects of our approach include: (1) constructing evaluation sets that exclude variants in the training sets of tools; for meta-predictors, even those from the training sets of the constituent tools must be excluded, (2) carefully inspecting whether the precomputed scores for a particular tool suffer from issues arising from missing data, outdated versions, and identifier mapping discrepancies and, if so, using a tool directly rather than precomputed scores deposited elsewhere, (3) calibrating tool scores and/or its assessment to account for the prior probability of pathogenicity that must be estimated for each reference set of interest, (4) reporting local posterior probabilities or local likelihood ratios, which provide additional information to standard evaluation metrics used in machine learning, and (5) providing thresholds for clinical use of a tool and the highest strength of evidence that it can provide. Laboratories using this strategy should choose one method and consistently use that method in evaluating all genes and variants at the determined evidential strength, rather than “cherry picking” amongst multiple methods. We nevertheless endorse gene-specific or gene domain-specific evaluations to identify evidence that demonstrates the superiority of a given tool that may be different from the general recommendations that we have specified (such as those in ClinGen VCEP recommendations).

### Implications for combining rules

Within the ACMG/AMP variant classification guidelines, there are several evidence criteria that can be applied independently of the PP3/BP4 computational tools, but whose underlying data may partially be captured by them, especially by the meta-predictor tools. The most obvious fall into two groups: allele frequency related codes, i.e., PM2 (absent from controls) and BS1 (more frequent than expected for disorder); and the key domain or critical residue(s) codes PM1 (located in functional domain or mutation hot spot), and PS1/PM5 (located at an amino acid where pathogenic variants have been seen). Increasing the strength of the PP3/BS4 computational tool codes, while at the same time including these other codes in the final classification, poses a risk of double-counting shared underlying attributes of these criteria and over-estimating the strength of evidence for or against pathogenicity.

There are reasonable approaches that can reduce double counting. For allele frequency data, classification should use *in silico* tools that do not make direct use of allele frequency, e.g., REVEL or BayesDel with the allele frequency option turned off (as was done in this study). Therefore, we recommend that the PM2 and BS1 codes may be combined with the PP3/BP4 codes at the strengths we have recommended here without any limits or additional criteria. However, the overlap of key domain / critical residue codes is more difficult to separate because it is oftentimes very highly correlated with attributes measured by an *in silico* tool (e.g., evolutionary conservation). Furthermore, it is challenging to separate these shared attributes for tools such as MutPred2 and VEST4 that implicitly incorporate some notion of structural and functional importance to each variant position. To address this potential overlap or double-counting of PP3/BP4 and PM1, we recommend that laboratories limit the sum of the evidence strength of PP3 and PM1 to Strong. This would allow PP3 to be invoked as Supporting or Moderate along with PM1 to be invoked as Moderate, which would be the same as limiting the sum of PP3 and BP4 to 4 points in the Bayes points implementation. Future stratified analyses, or integrated predictors amalgamating multiple codes, will be required to determine if or when these codes can be combined to provide even stronger evidence and the appropriate maximum allowable points for the different combinations.

### Limitations and future directions

There are caveats to our evaluation framework and, as a consequence, our recommendations. In this study, we estimated prior probabilities, calibrated tools, and estimated score thresholds using a genome-wide set of known disease-associated genes. As we noted above, in some circumstances it may be appropriate for a laboratory that focuses on a single or a few genes, to independently calibrate one of these tools using the method we describe here, which could lead to distinct numerical thresholds for the various evidence levels for that (those) specific gene(s). As well, VCEPs that assess specific genes can use our approach to establish predictive thresholds that can optimize the performance of computational tools in their specific systems.

Determination of different thresholds for the use of computational tools merits investigation of variants in ClinVar and potential re-classification of those where prediction models with insufficiently high (or low) scores played deciding roles. At the same time, the use of our approach will require detailed cataloguing of information in ClinVar such as the exact version of the tool, the raw prediction score, as well as whether a standalone tool or precomputed scores were used. This will be necessary to avoid circularities in future evaluations of computational tools. If our approach is adopted, we also suggest periodic investigation of the accuracy of the calibration we have proposed as ever-increasing data sets offer future potential to further improve the precision and accuracy of our thresholds.

Finally, it is important to emphasize that the approach presented herein is intended for the use alongside the ACMG/AMP rules^2^ and could lead to unintended consequences if used for variant classification outside of this setting.

## CONSORTIA

The ClinGen Consortium Sequence Variant Interpretation Working Group members include Leslie G. Biesecker, Steven M. Harrison (co-chairs), Ahmad A. Tayoun, Jonathan S. Berg, Steven E. Brenner, Garry R. Cutting, Sian Ellard, Marc S. Greenblatt, Peter Kang, Izabela Karbassi, Rachel Karchin, Jessica Mester, Anne O’Donnell-Luria, Tina Pesaran, Sharon E. Plon, Heidi L. Rehm, Natasha T. Strande, Sean V. Tavtigian, and Scott Topper.

## ACKNOWLEDGMENTS

We thank Drs. Joseph Rothstein and Weiva Sieh for filtering out variants in our set that were present in REVEL’s and constituent tools’ training sets. We thank Drs. Panagiotis Katsonis and Olivier Lichtarge for generating prediction scores for the Evolutionary Action approach. We also thank Drs. John Moult and Shantanu Jain for productive discussions. VP was supported by NIH grant K99LM012992. ABB and SMH were supported by NIH grant U24HG006834. AODL was supported by NIH grants U24HG011450, U01HG011755, and UM1HG008900. SVT was supported by NIH grants R01CA121245 and R01CA264971. MSG was supported by NIH grant U24CA258119. LGB was supported by NIH grant ZIAHG200359. PR and SEB were supported by NIH grant U24HG007346. PR was also supported by NIH grant U01HG012022. SEB was also supported by NIH grants U41HG009649 and R13HG006650, and a research agreement with Tata Consultancy Services. ClinGen is primarily funded by the National Human Genome Research Institute (NHGRI) with co-funding from the National Cancer Institute (NCI), through the following grants: Baylor/Stanford - U24HG009649, Broad/Geisinger - U24HG006834, and UNC/Kaiser - U24HG009650. The authors thank Julia Fekecs of NHGRI for graphics support. The content is solely the responsibility of the authors and does not necessarily represent the official views of the National Institutes of Health.

## DATA AND CODE AVAILABILITY

The three data sets, intermediate result files and code to calculate local posterior probabilities, estimate thresholds and plot figures in the paper are available here: https://github.com/vpejaver/clingen-svi-comp_calibration

