## Supplemental Materials for "Evidence-based calibration of computational tools for missense variant pathogenicity classification and ClinGen recommendations for clinical use of PP3/BP4 criteria"

### SUPPLEMENTAL METHODS

#### Alternative strategies for interval definition

We also investigated two other strategies to define intervals corresponding to the relevant evidential support. The first strategy used the global likelihood ratio and defined the threshold for the supporting level of evidence as

$$\tau_{\mathrm{Su}}^{\mathrm{P}} = \min\{\tau : \forall t \geq \tau, \mathrm{LR}^{+}(t, \infty) \geq 2.406\},$$

where $\mathrm{LR}^{+}(\tau, \infty)$ is the positive likelihood ratio obtained when predictions $s \in [\tau, \infty)$ are considered pathogenic and used to compute the posterior odds of pathogenicity using equations 1 and 2 in the main text. The remaining thresholds from $\mathcal{T}_{\mathrm{P}}$ were defined as in the equation above except that the likelihood ratio levels were selected from Table 1. The same procedure was repeated for the benignity set $\mathcal{T}_{\mathrm{B}}$ using the negative likelihood ratio to define levels of evidential support. All intervals $\mathrm{I}^{\mathrm{P}}(\text{evidence level})$ and $\mathrm{I}^{\mathrm{B}}(\text{evidence level})$, where $\text{evidence level} \in \{\mathrm{Su}, \mathrm{Mo}, \mathrm{St}, \mathrm{VSt}\}$, were therefore established from the threshold sets $\mathcal{T}_{\mathrm{P}}$ and $\mathcal{T}_{\mathrm{B}}$.

The second strategy for selecting threshold sets defined all thresholds simultaneously by satisfying $\mathrm{LR}^{+}(\tau_{\mathrm{Su}}^{\mathrm{P}}, \tau_{\mathrm{Mo}}^{\mathrm{P}}) \geq 2.406$, $\mathrm{LR}^{+}(\tau_{\mathrm{Mo}}^{\mathrm{P}}, \tau_{\mathrm{St}}^{\mathrm{P}}) \geq 5.790$, $\mathrm{LR}^{+}(\tau_{\mathrm{St}}^{\mathrm{P}}, \tau_{\mathrm{VSt}}^{\mathrm{P}}) \geq 33.53$, and $\mathrm{LR}^{+}(\tau_{\mathrm{VSt}}^{\mathrm{P}}, \infty) \geq 1124.000$, where $\mathrm{LR}^{+}(\tau_{\mathrm{Su}}^{\mathrm{P}}, \tau_{\mathrm{Mo}}^{\mathrm{P}})$ was obtained when predictions $s \in [\tau_{\mathrm{Su}}^{\mathrm{P}}, \tau_{\mathrm{Mo}}^{\mathrm{P}})$ were considered pathogenic and used to compute the posterior odds of pathogenicity. Since this approach may not have a unique solution, a greedy approach was used to optimize threshold intervals.

#### Suggested modification to Tavtigian et al. framework

To use BP4_Moderate with other evidence we propose a modification to Equation 2 in Tavtigian et al.:

$$OP = O_{PVSt}^{-\frac{N_{BSu}}{X^3} - \frac{N_{BMo}}{X^2} - \frac{N_{BSt}}{X}}$$

where $OP$ is the Odds of pathogenicity, $O_{PVSt}$ is the Odds of pathogenicity corresponding to Very Strong evidence for pathogenicity, $N$ is the number of lines of evidence for benignity (with the subscript indicating the strength levels) and $X$ is a scaling factor. While this equation is presented this way here to preserve the notation of Tavtigian et al., there is a one-to-one correspondence between this equation and Equation 5 of this study. $OP$ corresponds to the positive likelihood ratio ($\mathrm{LR}^{+}$), $O_{PVSt}$ corresponds to $c$, $N$ is corresponds to $n$, and $X$ is set to 2 (as in the original Tavtigian et al. framework).

### SUPPLEMENTAL TABLES

**Table S1. Estimated thresholds for all tools in this study corresponding to the four pathogenic and four benign intervals.** The Confidence Interval (CI) column indicates the one-sided 95% confidence bound. For the selection of thresholds, the confidence bounds were chosen (except for FATHMM and SIFT, these would be higher than the point estimates for PP3 and these would be lower than the point estimates for BP4). In this manner, the recommended thresholds were more stringent and accounted for uncertainty to the best extent possible. A "–" implies that the given tool did not meet the posterior probability (likelihood ratio) threshold.

| Method | PP3_VeryStrong | | PP3_Strong | | PP3_Moderate | | PP3_Supporting | |
|---|---|---|---|---|---|---|---|---|
| | Estimate | CI | Estimate | CI | Estimate | CI | Estimate | CI |
| BayesDel | - | - | 0.49 | 0.50 | 0.24 | 0.27 | 0.11 | 0.13 |
| CADD | - | - | - | - | 26.7 | 28.1 | 25.0 | 25.3 |
| EA1.0 | - | - | 0.981 | - | 0.787 | 0.821 | 0.628 | 0.685 |
| FATHMM | - | - | - | - | -4.79 | -5.04 | -4.05 | -4.14 |
| GERP++ | - | - | - | - | - | - | - | - |
| MPC | - | - | - | - | 1.735 | 1.828 | 1.314 | 1.36 |
| MutPred2 | - | - | 0.924 | 0.932 | 0.793 | 0.829 | 0.683 | 0.737 |
| PhyloP | - | - | - | - | 9.664 | 9.741 | 7.085 | 7.367 |
| PolyPhen-2 | - | - | - | - | 0.998 | 0.999 | 0.97 | 0.978 |
| PrimateAI | - | - | - | - | 0.844 | 0.867 | 0.766 | 0.790 |
| REVEL | - | - | 0.918 | 0.932 | 0.736 | 0.773 | 0.629 | 0.644 |
| SIFT | - | - | - | - | 0.000 | 0.000 | 0.002 | 0.001 |
| VEST4 | - | - | 0.958 | 0.965 | 0.838 | 0.861 | 0.747 | 0.764 |
| **Method** | **BP4_Supporting** | | **BP4_Moderate** | | **BP4_Strong** | | **BP4_VeryStrong** | |
| | **Estimate** | **CI** | **Estimate** | **CI** | **Estimate** | **CI** | **Estimate** | **CI** |
| BayesDel | -0.16 | -0.18 | -0.27 | -0.36 | -0.54 | - | - | - |
| CADD | 23.0 | 22.7 | 20.4 | 17.33 | 1.898 | 0.154 | - | - |
| EA1.0 | 0.286 | 0.262 | 0.149 | 0.069 | - | - | - | - |
| FATHMM | 2.20 | 3.32 | 4.12 | 4.69 | - | - | - | - |
| GERP++ | 3.11 | 2.70 | 1.36 | -4.54 | - | - | - | - |
| MPC | - | - | - | - | - | - | - | - |
| MutPred2 | 0.408 | 0.391 | 0.208 | 0.197 | 0.023 | 0.01 | - | - |
| PhlyoP | 2.054 | 1.879 | 0.323 | 0.021 | - | - | - | - |
| PolyPhen-2 | 0.158 | 0.113 | 0.025 | 0.009 | - | - | - | - |
| PrimateAI | 0.541 | 0.483 | 0.393 | 0.362 | - | - | - | - |
| REVEL | 0.348 | 0.290 | 0.238 | 0.183 | 0.046 | 0.016 | 0.003 | 0.003 |
| SIFT | 0.061 | 0.08 | 0.235 | 0.327 | - | - | - | - |
| VEST4 | 0.474 | 0.449 | 0.325 | 0.302 | 0.073 | - | - | - |

**Table S2. Percentage of missing predictions for all tools for the three data sets in this study.**

| Method | ClinVar 2019 | gnomAD | ClinVar 2020 |
|---|---|---|---|
| BayesDel | 0.0 | 0.0 | 0.0 |
| CADD | 0.6 | 0.0 | 0.0 |
| EA1.0 | 16.1 | 11.0 | 10.3 |
| FATHMM | 14.0 | 13.4 | 13.1 |
| GERP++ | 2.3 | 1.5 | 1.7 |
| MPC | 29.5 | 23.5 | 25.5 |
| MutPred2 | 6.5 | 0.0 | 0.6 |
| PhyloP | 2.3 | 1.5 | 1.7 |
| PolyPhen-2 | 19.4 | 16.7 | 14.9 |
| PrimateAI | 8.7 | 5.3 | 5.7 |
| REVEL | 4.0 | 2.2 | 2.5 |
| SIFT | 21.5 | 15.3 | 15.1 |
| VEST4 | 8.8 | 6.6 | 6.8 |

### SUPPLEMENTAL FIGURES

| | BayesDel | CADD | EA | FATHMM | GERP++ | MPC | MutPred2 | PhyloP | PolyPhen-2 | PrimateAI | REVEL | SIFT | VEST4 |
|---|---|---|---|---|---|---|---|---|---|---|---|---|---|
| BayesDel | 1.00 | | | | | | | | | | | | |
| CADD | 0.68 | 1.00 | | | | | | | | | | | |
| EA | 0.63 | 0.56 | 1.00 | | | | | | | | | | |
| FATHMM | 0.55 | 0.13 | 0.09 | 1.00 | | | | | | | | | |
| GERP++ | 0.50 | 0.78 | 0.35 | 0.08 | 1.00 | | | | | | | | |
| MPC | 0.45 | 0.43 | 0.33 | 0.14 | 0.28 | 1.00 | | | | | | | |
| MutPred2 | 0.74 | 0.61 | 0.62 | 0.23 | 0.42 | 0.47 | 1.00 | | | | | | |
| PhyloP | 0.64 | 0.70 | 0.48 | 0.12 | 0.67 | 0.40 | 0.59 | 1.00 | | | | | |
| PolyPhen-2 | 0.64 | 0.67 | 0.65 | 0.11 | 0.39 | 0.43 | 0.60 | 0.52 | 1.00 | | | | |
| PrimateAI | 0.63 | 0.64 | 0.43 | 0.17 | 0.49 | 0.55 | 0.64 | 0.63 | 0.50 | 1.00 | | | |
| REVEL | 0.89 | 0.58 | 0.61 | 0.58 | 0.38 | 0.43 | 0.73 | 0.59 | 0.64 | 0.55 | 1.00 | | |
| SIFT | 0.42 | 0.49 | 0.46 | 0.07 | 0.30 | 0.25 | 0.36 | 0.32 | 0.47 | 0.29 | 0.37 | 1.00 | |
| VEST4 | 0.80 | 0.70 | 0.66 | 0.17 | 0.49 | 0.45 | 0.77 | 0.69 | 0.69 | 0.68 | 0.72 | 0.43 | 1.00 |

**Figure S1. Pairwise correlations among all methods on the *gnomAD* data set.** The Pearson correlation coefficients are shown for all pairs of tools. Darker squares indicate higher coefficients. Note that the figure accounts for the fact that SIFT and FATHMM output scores in an inverted scale (higher scores indicating benignity).
